# Mapping the interaction sites of Influenza A viruses and human complement Factor H

**DOI:** 10.1101/2023.09.15.557969

**Authors:** Iman Rabeeah, Elizabeth Billington, Beatrice Nal-Rogier, Jean-Remy Sadeyen, Ansar Pathan, Munir Iqbal, Nigel Temperton, Peter F. Zipfel, Christine Skerka, Uday Kishore, Holly Shelton

## Abstract

The complement system is an innate immune mechanism against microbial infection. It involves a cascade of effector molecules that is activated via classical, lectin and alternative pathways. Consequently, many pathogens bind to or incorporate in their structures host negative regulators of the complement pathways as an evasion mechanism. Factor H (FH) is a negative regulator of the complement alternative pathway that protects “self” cells of the host from non-specific complement attack. Viruses including human influenza A viruses (IAVs) have been shown to bind to FH. Here we show that IAVs of both human and avian origin can bind directly to human FH and the interaction is mediated via the IAV surface glycoprotein haemagglutinin (HA). HA bound to common pathogen binding footprints on the FH structure, complement control protein modules, CCP 5-7 and CCP 15-20. The FH binding to H1 and H3 showed that the interaction overlapped with the receptor binding site of both HAs but the footprint was more extensive for the H3 HA than the H1 HA. The HA - FH interaction impeded the initial entry of H1N1 and H3N2 IAV strains but its impact on viral multicycle replication in human lung cells was strain specific. The H3N2 virus binding to cells was significantly inhibited by preincubation with FH, whereas there was no alteration in replicative rate and progeny virus release for human H1N1 or avian H9N2 and H5N3 IAV strains. We have mapped the interaction between IAV and FH, the significance of which for the virus or host is yet to be elucidated.

## Introduction

Influenza A viruses (IAVs) are enveloped viruses that carry a segmented, negative sense, single-stranded RNA genome. Human H1N1 and H3N2 IAVs cause regular seasonal outbreaks in the human population where the disease can result in severe clinical outcomes in children, elderly, pregnant women, and immune compromised individuals. The public health burden of these annual outbreaks is significant with up to 650,000 excess deaths estimated annually by the WHO from influenza infections [1]. In addition, a vast array of different IAV subtypes circulate in the natural reservoir population of wild aquatic birds and sporadically these avian IAVs can cross the host species barrier and result in infection of humans, often with high case fatality rates, such as H7N9 or H5N1 [2–4].

The complement system is one of the very first innate immune responses mounted by vertebrates against infection by pathogens. The complement system is activated using three different pathways; classical, lectin and alternative; it can detect pathogens and trigger clearance mechanisms via virolysis, prom-inflammatory cytokine production and potentiation of the adaptive immunity [5, 6]. It has been demonstrated that the complement system can control IAV infections via the classical pathway, whereby virus-specific or non-specific natural IgG antibodies bind to the virus allowing its association with C1 of the classical pathway [7, 8]. The subsequent deposition of complement components onto the virus surface leads to virus aggregation and inhibition of receptor binding and thus neutralisation [9, 10]. The lectin pathway acting through the binding of either mannan-binding lectin (MBL) or L-ficolin to IAV surface Haemagglutinin (HA) protein and the alternative pathway have also both been demonstrated to contribute to the protection of the host from IAV [11, 12] .

In order to successfully replicate and transmit, many viruses have evolved complement evasion strategies [13]. Incorporation of complement regulators into virion structures, such as the central complement inhibitors CD55 and CD59 which are found in the HIV membrane envelope is one strategy [14]. Virus mimicry of complement regulators also occurs, for example, the gC of Herpes Simplex Virus 1 mimics the action of CD55 or CD59, inhibiting the C3b molecule central to the complement activation [15, 16]. Additionally, viruses have been shown to perturb complement protein expression levels. In Hepatitis C virus infected cells, there is a down-regulation of C2, C4, C3 and C9 expression which results in reduced complement activation [17–19]. IAV is known to incorporate CD59 into its envelope in order to protect against membrane attack complex lysis by preventing C9 polymerisation [20]. In addition, the Matrix (M1) protein of IAV binds to the recognition subcomponent C1q which prevents classical pathway activation. However, at what point in the virus lifecycle the M1 protein, which is internal to the virus structure, gets to contact C1q is not known [21].

Factor H (FH) is a large, 150kDa plasma glycoprotein, which acts as a negative regulator of the alternative pathway [22]. The alternative pathway, unlike the classical and lectin pathways, does not require a specific binding event to get activated, but instead, is initiated by spontaneous hydrolysis of C3. FH regulates this process by binding to C3b which inhibits the formation of the critical C3 convertase enzyme. FH is also a co-factor for factor I, a protease that cleaves C3b into an inactive form (iC3b). These actions by FH act to protect the hosts’ self-structures from this non-directed alternative pathway [23]. Several pathogens bind to FH in the host and use this as an evasion mechanism to protect against alternative pathway mediated lysis [24, 25]. Pathogenic bacteria including *Borrelia burgdorferi*, *Staphylococcus aureus*, *Streptococcus pneumoniae, Neisseria meningitidis* and *Neisseria gonorrhoeae* encode proteins that bind to FH, thus providing mimicry of the hosts self surfaces which leads to increased survival of the pathogens [24].

Viruses can also directly bind FH to evade the effect of complement. The NS1 protein of the Flavivirus, West Nile Virus (WNV), has been shown to directly interact with human FH [26]. The WNV NS1 protein is a secreted glycoprotein which can then bind back to the surface of cells, so its interaction with FH may reduce the recognition of infected cells by the complement leading to pathogen survival. Zaiss et al. have demonstrated that the capsid proteins of Adeno-associated virus (AAV) can also directly interact with human FH and this allows the inhibition of C3 convertase on the virion surface [27]. In addition, human IAV strains have been shown to interact directly with FH [28].

In this study, we confirm that there is a direct interaction between human and avian subtypes of IAVs and FH purified from human serum. We demonstrate that human FH binds to the IAV surface HA glycoprotein using its common pathogen binding domains, which are also involved in FH binding to bacterial proteins. The interaction site of FH on the HA of human H1 and H3 subtypes is in the vicinity of the receptor binding site and the consequence of this is an initial inhibition of viral entry to human cells when FH is present. However, there is a more extensive FH binding footprint on the H3 HA compared to the H1 HA and this results in modulation of multicycle replication that is IAV strain dependent.

## Materials and Methods

### Cells

A549 and MDCK cells (Cell Culture Central Services Unit, The Pirbright Institute) were grown in Complete Medium, DMEM (Gibco-Invitrogen, Inc.) supplemented with 10% (v/v) foetal bovine serum (FBS) (Biosera, Inc.), 1% (v/v) penicillin/streptomycin (Sigma-Aldrich) and 1% (v/v) non-essential amino acids (Sigma-Aldrich), and maintained at 37°C in 5% CO_2_.

### Viruses

The various Influenza A viruses used in this study are listed in Table 1.

**Table 1:** Viruses used in this study.

| IAV strain | Subtype | Origin |
| --- | --- | --- |
| A/Hong Kong/1774/99 | H3N2 | Leo Poon, University of Hong Kong |
| A/Hong Kong/4801/14 | H3N2 | NIBSC (NYMC-X-263) |
| A/England/195/09 | H1N1 | Wendy Barclay, Imperial College London [35] |
| A/Michigan/45/2015 | H1N1 | NIBSC (NYMC X-275) |
| A/chicken/Pakistan/UDL01/08 | H9N2 | Reverse genetics [36] |
| A/duck/Singapore/3/97 | H5N3 | Munir Iqbal, Pirbright Institute |

Human IAV strains were propagated in MDCK cells in the presence of 1 µg/ ml TPCK trypsin (Sigma) and harvested after 72 hours. Avian influenza strains were propagated in specific pathogen free (SPF) 10-day old embryonated chicken eggs (VALO BioMedia, Germany), the allantoic fluid harvested after 48 hours. Purification of IAVs was performed by ultra-centrifugation on to a 30% sucrose cushion. Briefly, virus solutions were clarified of debris by centrifugation at 10,000 rpm at 4°C for 30 minutes in a chilled SW32Ti rotor (Beckman Coulter), followed by pelleting through a 30% cold sucrose solution at 25,000 rpm for 90 minutes. The viral pellet was recovered and resuspended in PBS. Viral titre was determined by plaque assay on MDCK cells, as previously described [36], or by TCID50 assay. Briefly, TCID50 was performed by diluting virus in half Log_10_ series across a 96 well tissue culture plate in serum free DMEM. 5 x 10_5_ MDCK cells per ml were prepared in serum free DMEM containing 2% v/v penicillin/streptomycin (Sigma-Aldrich) and 2 µg/ml TPCK Trypsin (Sigma). 100 µl of MDCK cells were added to each well and incubated at 37°C with 5% CO_2_ for 3 days. Cells were fixed with ice cold 50:50 methanol:acetone for 20 minutes and stained with 0.1% crystal violet solution containing methanol. Cytopathic effect (CPE) was recorded for each well and TCID50 calculated by the Reed-Muench method [37]

### Lentiviral pseudotyped particles

MLV particles carrying a firefly luciferase expression plasmid were pseudotyped with IAV HA alone or IAV HA and NA, the VSV-G protein or in the absence of an envelope protein. Pseudotyped lentivirus particles were produced in HEK293T cells as earlier described [38, 39]. Pseudotyped lentivirus particles were titrated by quantification of relative light units produced by firefly luciferase activity in transduced HEK293T cells (Table 2).

**Table 2:** Envelope proteins used to produce pseudotyped lentiviral particles.

| Viral strain | IAV Subtype | Envelope receptors |
| --- | --- | --- |
| A/Udorn/307/1972 | H3N2 | HA |
| A/California/7/2004 | H3N2 | HA |
| A/Texas/1/1977 | H3N2 | HA |
| A/Solomon Island/3/2006 | H1N1 | HA |
| A/England/195/2009 | H1N1 | HA |
| A/Hong Kong/33982/2009 | H9N2 | HA |
| A/FPV/Rostock/1934 | H7N1 | HA |
| A/Shanghai/02/2013 | H7N9 | HA |
| A/Vietnam/1194/2004 | H5N8 | HA |
| A/Vietnam/1194/2004 | H5N8 | HA+NA |
| De-envelope protein |  | - |
| Vesicular stomatitis virus |  | G |

### Purification of FH protein

Human FH was purified from pooled human plasma (TCS Biosciences LTD) via a three-step affinity chromatography protocol [40]. Step 1 involved removal of plasmin/plasminogen from plasma. A L-Lysine – Sepharose column [1 g of Cyanogen bromide (CNBr)-activated Sepharose 4B (Sigma)] was added to 15 ml of 100 mM NaHCO3 (Sigma) buffer, pH 8.9 with 2 g of L-lysine monohydrochloride (Fisher Scientific). The column was equilibrated with binding buffer [100 mM sodium phosphate (Sigma), 150 mM NaCl (Sigma), 15 mM EDTA (Sigma), pH 7.4)] and 50 ml of human plasma applied at a rate of 1 ml/minute. The column was washed with 100 ml of binding buffer and the flow-through was collected and then dialysed against buffer containing 25 mM Tris–HCL (Sigma), 140 mM NaC1, 0.5 mM EDTA, pH 7.4.

Step 2 involved removal of IgG binding proteins from plasma. 20 mg of purified human IgG in PBS was immobilised to CNBr-activated Sepharose (Sigma) (10 mg IgG/ml packed Sepharose). Reactive sites in the Sepharose were blocked by the addition of 20 volumes of 100 mM Tris-HCl, 150 mM NaCl, pH 8.5 for 2 hours. The column was equilibrated in running buffer (25 mM Tris–HCl, 140 mM NaCl, 0.5 mM EDTA, pH 7.4). Plasmin/ Plasminogen free human plasma was then passed through the IgG– Sepharose column in running buffer and the column washed with 100 ml of running buffer. The flow-through was again collected.

Step 3, the last chromatographic step, involved use of anti-human FH (OX23) monoclonal antibody [41]. 8 mg of purified anti-FH OX23 antibody (MRC Immunochemistry Unit, Oxford) from mouse hybridoma cell line culture supernatant was immobilised on CNBr-activated Sepharose (Sigma) (2 mg/ml packed Sepharose). The column was equilibrated in running buffer and then plasma depleted of plasmin/plasminogen and IgG binding proteins was passed through the antibody affinity column. Column was washed extensively with 100 ml of running buffer and then FH protein was eluted in 1 ml fractions by 3 M MgCl2. Fractions containing FH were pooled and dialysed against PBS, and stored at -20°C in aliquots. The purity of FH was assessed by 10% w/v SDS-PAGE (Supplementary Figure 1A).

### HA peptides

H1 and H3 peptides were custom made by Alta Bioscience Ltd, based on the sequence of H1N1 A/England/195/09 and H3N2 A/Hong Kong/1774/99. Both HA sequences were covered by 56 peptides; 55 of the peptides for each HA were 20 amino acid long with an overlap along the protein sequence of 10 amino acids; the final peptide sequences were 16 (H1) or 15 (H3) amino acids in length. Each peptide had an N-terminal biotin tag and an amide at the C-terminus. The sequences of the synthetic peptides are listed in Supplementary Table 1.

### ELISA

The interaction of IAV and pseudotyped lentivirus with FH or FH CCP fragments was assessed by ELISA. 10^4^ purified IAV or 10^4^ relative light units (RLU) of pseudotype lentivirus were adsorbed onto 96-well Maxisorp microtitre plates (Fisher Scientific) in carbonate-bicarbonate (CBC) buffer pH 9.6 (Sigma) overnight at 4°C. Plates were then washed with PBST [PBS with 0.1% Tween-20 (Sigma)]. Wells were blocked for 1 hour at room temperature with 5% w/v BSA in PBS (blocking buffer). Purified FH or FH CCP fragments diluted in PBS were added at various known concentrations and incubated for 2 hours at room temperature. Wells were washed with PBST, and primary antibody, either OX24 monoclonal anti-FH (purified from mouse hybridoma supernatant; MRC Immunochemistry Unit, Oxford) or goat anti-human FH (Complement Technology) diluted in blocking buffer were applied for 1 hour at room temperature. After washing in PBST, secondary antibody, rabbit anti-mouse-HRP (DAKO) or rabbit anti-goat (Abcam), diluted in blocking buffer, was applied for 1 hour at room temperature. Following incubation, wells were washed in PBST and the colour was developed using 3,3′,5,5′-Tetramethylbenzidine (TMB) substrate (Sigma). The reaction was stopped using 2 N H_2_SO_4_ (Sigma). The absorbance was read at 450 nm using Absorbance microplate Reader ELX 808 (Bio-Tek).

### Virus cell entry assay

H1N1 A/England/195/09 or H3N2 A/Hong Kong/1774/99/ IAV at a MOI of 1 was preincubated in the presence or absence of 100 µg/ml of purified FH for 1 hour a 37°C. Virus was then inoculated onto A549 cells and allowed to infect for 1 hour at 37°C before being removed. Cells were then incubated at 37°C with 5% CO_2_ in serum free DMEM containing 2% penicillin/streptomycin (Sigma) and 1 µg/ml TPCK Trypsin (Sigma) for either 4 hours or 8 hours. At the appropriate timepoint, cells were either fixed with ice cold 50:50 methanol: acetone for 5 minutes at room temperature, or lysed by scraping into buffer containing 1% Triton X 100, 20 mM Tris-HCl, 150 mM NaCl, 1 mM EDTA, together with 1% anti-proteases cocktail (Themo Scientific). Fixed cells were immune stained for viral NP protein using mouse anti-NP (provided by Munir Iqbal, Pirbright Institute) and goat anti-mouse HRP conjugate, followed by development with AEC Substrate Chromogen (DAKO); images were acquired via EVOS XL Core Cell Imaging system (Invitrogen). Lysed cells were run on a 10% SDS-PAGE under reducing conditions and western blotting was carried out. The blot was probed using mouse monoclonal anti-M1 (Bio-Rad) and goat anti-mouse IRDye 800CW (LiCor) and visualised using the Odessey Imaging system (LiCor).

### Virus infection assay

A549 cells were infected with a MOI 0.1 of H1N1 (A/England/195/09) or H3N2 (A/Hong Kong/1174/99) IAV that had been preincubated with various concentrations of FH (0, 5, 50 or 100 µg). Alternatively, A549 cells were infected with a MOI 0.1 of H9N2 (A/chicken/Pakistan/UDL01/08) or H5N3 (A/duck/Singapore/3/97) IAV that had been preincubated with 100 µg of FH. Cells were then incubated at 37°C under 5% CO_2_ in serum free DMEM containing 2% penicillin/streptomycin (Sigma) and 1 µg/ml TPCK Trypsin (Sigma) for 24 hours, after which the cell supernatants were removed from infected A549 cells and titrated for infectious virus by TCID50 assay.

## Results

### Human and avian Influenza A viruses bind to human FH protein

To test the interaction between IAV and human FH, purified FH and IAV strains were produced (Supplementary Figure 1). Using an ELISA assay, we tested the binding of a range of concentrations (0 – 80 µg/well) of affinity purified human FH protein to 10^4^ pfu of human and avian IAV viruses (Figure 1). Two strains of human H1N1 virus (A/Michigan/45/2015 and A/England/195/2009) and two human H3N2 viruses (A/Hong Kong/1774/1999 and A/Hong Kong/4801/14) were tested. Significant binding (p < 0.05) was seen for all four human viruses at the FH concentration of 40 µg/well (Figure 1A-D). Control groups included the absence of FH (0 µg/well), absence of any IAV (No Virus), lentivirus particles with no envelope proteins (ENV) and lentivirus particles with VSV-G protein embedded in the membrane, the later three control groups had 80 µg/ well of FH applied to them. Since avian viruses can also sporadically cross the species barrier to infect humans, we also examined possible interaction between human FH and avian influenza viruses. Similar to the human IAV strains, FH protein (40µg/well) bound significantly to H9N2 (A/chicken/ Pakistan/UDL01/2008) as well as H5N3 (A/duck/Singapore/3/97) IAV (Figure 1 E, F).

**Figure 1.**
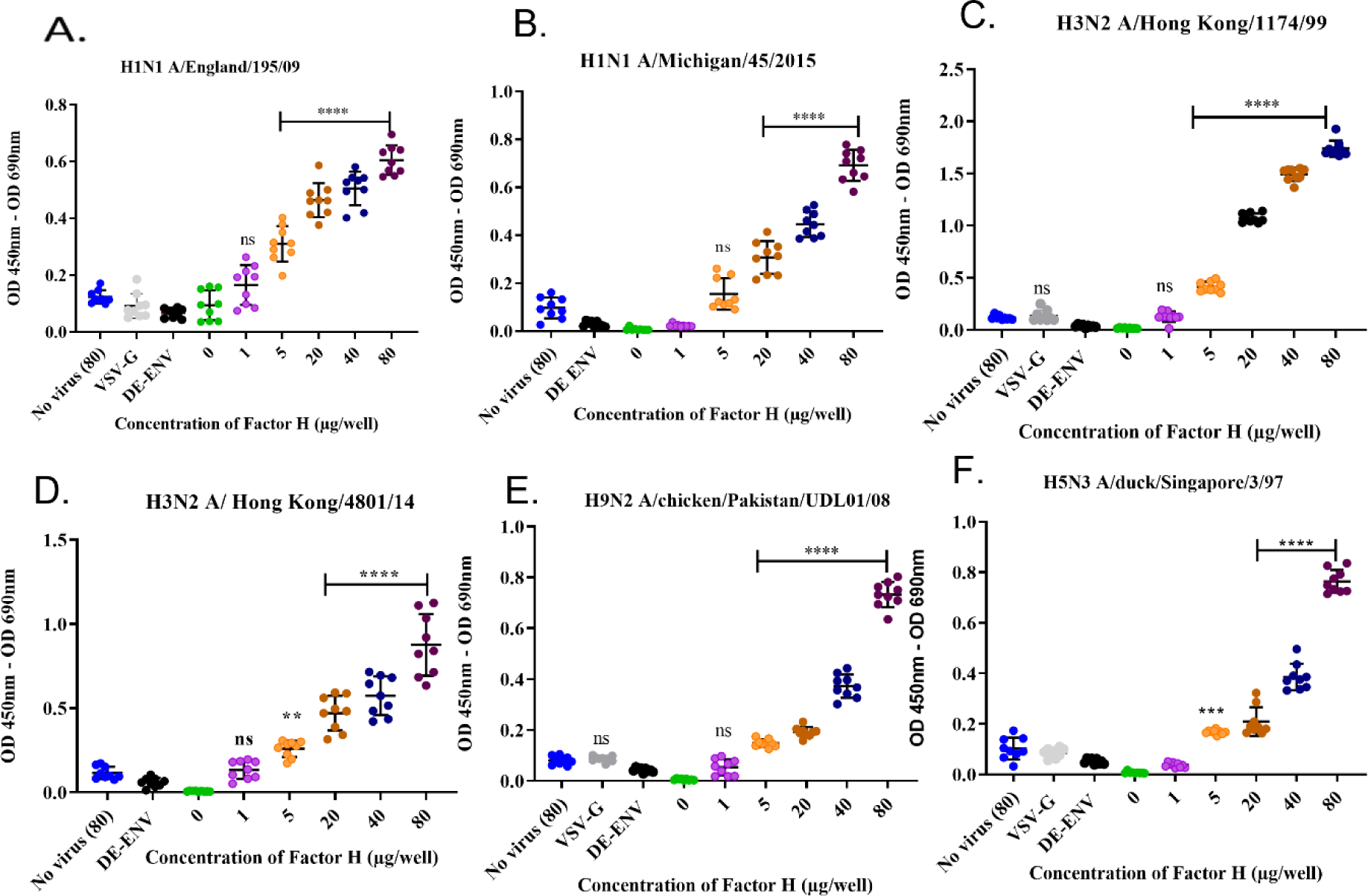
Influenza A viruses bind to human factor H protein. An ELISA was carried out to assess the direct interaction between IAV strains (A) H1N1 A/England/195/09, (B) A/Michigan/45/2015, (C) H3N2 A/Hong Kong/1174/99, (D) H3N2 A/Hong Kong/4801/14, (E) H9N2 A/chicken/Pakistan/UDL01/09 and (F) H5N3 A/duck/Singapore/3/97 and purified FH protein. Concentrations of 0, 1, 5, 20, 40 and 80 µg/well of purified FH were tested with 10^4^ pfu/well of IAV alongside the controls of 80 µg FH with no virus (No virus (80), lentivirus particles with VSV-G in the envelope (VSV-G) and lentivirus particles generated with no envelope protein (DE-ENV). FH binding was detected by anti-FH monoclonal antibody (OX24). Individual values from three independent experiments having three replicates per experiment are shown with mean OD 450nm – OD 690nm and error bars indicating standard deviation. Significant increased binding was calculated by One way ANOVA and post hoc Dunnetts multiple comparison test to the control group of no virus (80), ** p≤0.01, *** p ≤0.001 and **** p ≤0.0001.

### IAV binds human FH at common pathogen binding surfaces

A wide variety of pathogens including bacteria, fungi and viruses have been shown to interact with human FH protein, predominantly via its CCP5-7 and CCP19-20 modules. These CCP modules are also where the known binding sites for host cell ligands are located [29]. Binding to these sites does not interfere with FH interaction with C3b; thus, it does not functionally compromise the ability of FH to downregulate the complement alternative pathway [29]. Using recombinant fragments corresponding to different portions of human FH protein, we mapped the interaction of human and avian IAV strains using the ELISA (Figure 2A-D). All CCP fragments were detected by the polyclonal goat anti-human FH antibody used in ELISA (Supplementary figure 2A). We found that all four IAV strains bound to the FH CCP8-20 fragment with the greatest signal; significant binding was observed for the FH fragments encompassing CCP1-7 and CCP15-18. In addition, for the H9N2 virus, we found significant interaction with the CCP8-11 fragment but not for the H1N1, H3N2 or H5N3 strains. There was no significant binding to any of the IAV strains by FH fragments containing CCP1-4 or CCP11-15 which suggested that the footprint for IAV on the FH protein could be refined to CCP5-7 and CCP15-20 (Figure 2E).

**Figure 2.**
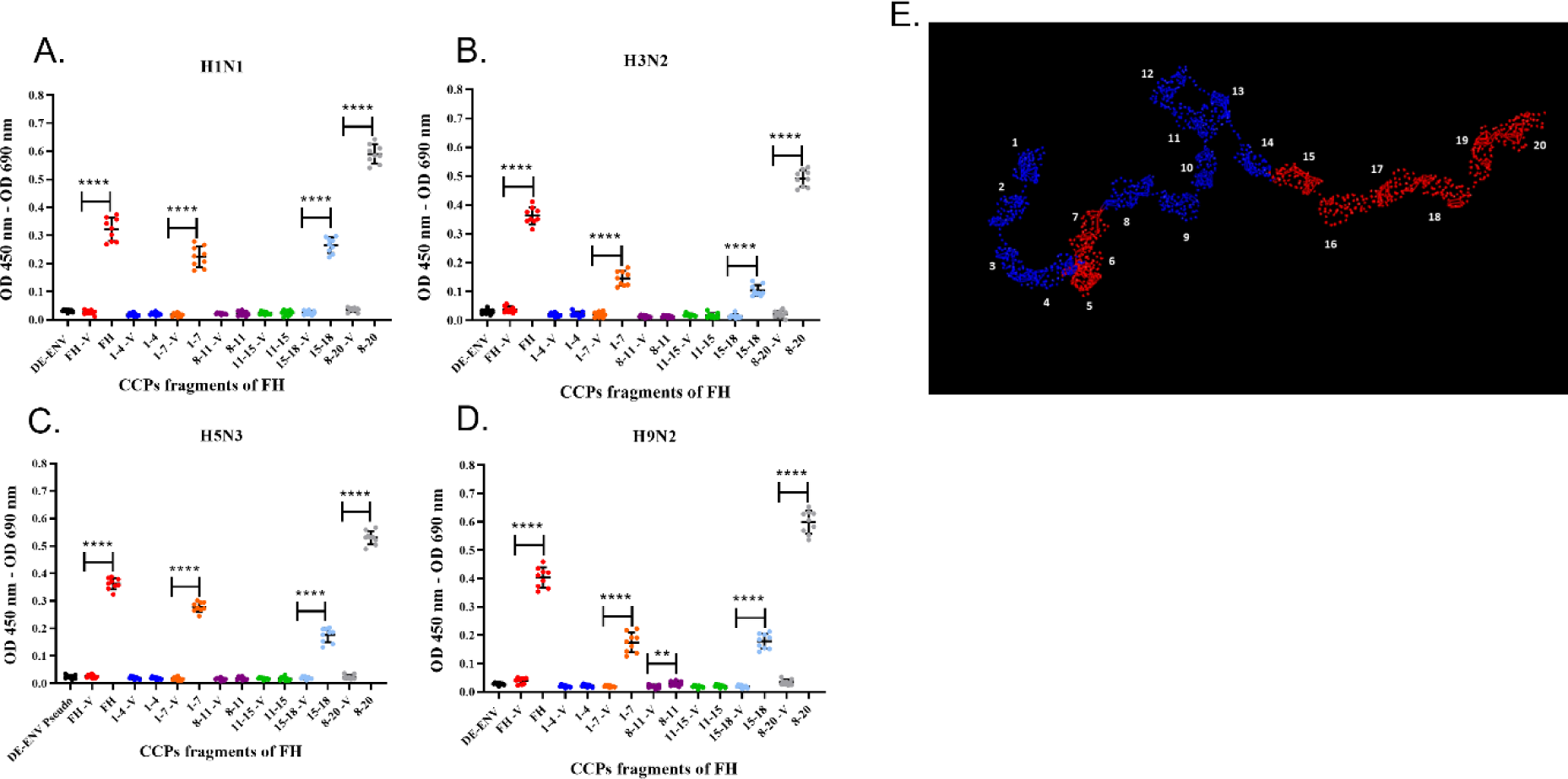
IAV strains bind human factor H at CCP domains 5-7 and 15-20. 10^3^ pfu/well of IAV; (A) H1N1 A/England/195/09, (B) H3N2 A/Hong Kong/1174/99, (C) H5N3 A/duck/Singapore/3/97 and (D) H9N2 A/chicken/Pakistan/UDL01/09 were used in an ELISA with 5 µg/well of purified FH protein (FH) or CCP fragments of FH, (CCP 1-4, 1-7, 8-11, 11-15, 15-18 and 8-20). Each fragment was assessed against its own controls of 5 µg FH with no virus (-V) and lentivirus particles generated with no envelope protein (DE-ENV). FH binding was detected by polyclonal goat anti-human FH (Complement Technology). Individual values from three independent experiments carrying three replicates per experiment are displayed with mean OD 450nm – OD 690nm and error bars indicating standard deviation. Significant binding was calculated by unpaired two-way T-test to the CCP fragment control group of -V, ** p≤0.01 and **** p ≤0.0001. (E) PyMOL generated structure from 1HAQ protein database entry of human FH, 20 CCP domains numbered and domains in red indicated predicted IAV interaction sites, CCPs 5-7 and CCPs 15-20 from ELISA data.

### IAV binding to human FH is mediated via the viral Haemagglutination (HA) protein

To confirm that the interaction between IAV and human FH was mediated via the viral surface glycoprotein Haemagglutinin (HA), we performed the ELISA using lentiviruses pseudotyped with HA protein of different IAV subtypes. These lentiviruses only express the IAV HA protein on the surface of the virion. We tested a panel of human H1 and H3 HA proteins and avian H9, H7 or H5 HA, all of which bound the FH significantly compared to the controls (Figure 3A), which was lentivirus pseudotyped with the vesicular stomata virus (VSV) surface protein G. In addition, we tested a lentivirus pseudotyped with both a H5 HA and N1 NA proteins from avian IAV and found that the addition of NA into the viral envelope did not significantly alter the binding of the FH to virus particle compared to the particle displaying H5 HA alone (Figure 3).

**Figure 3.**
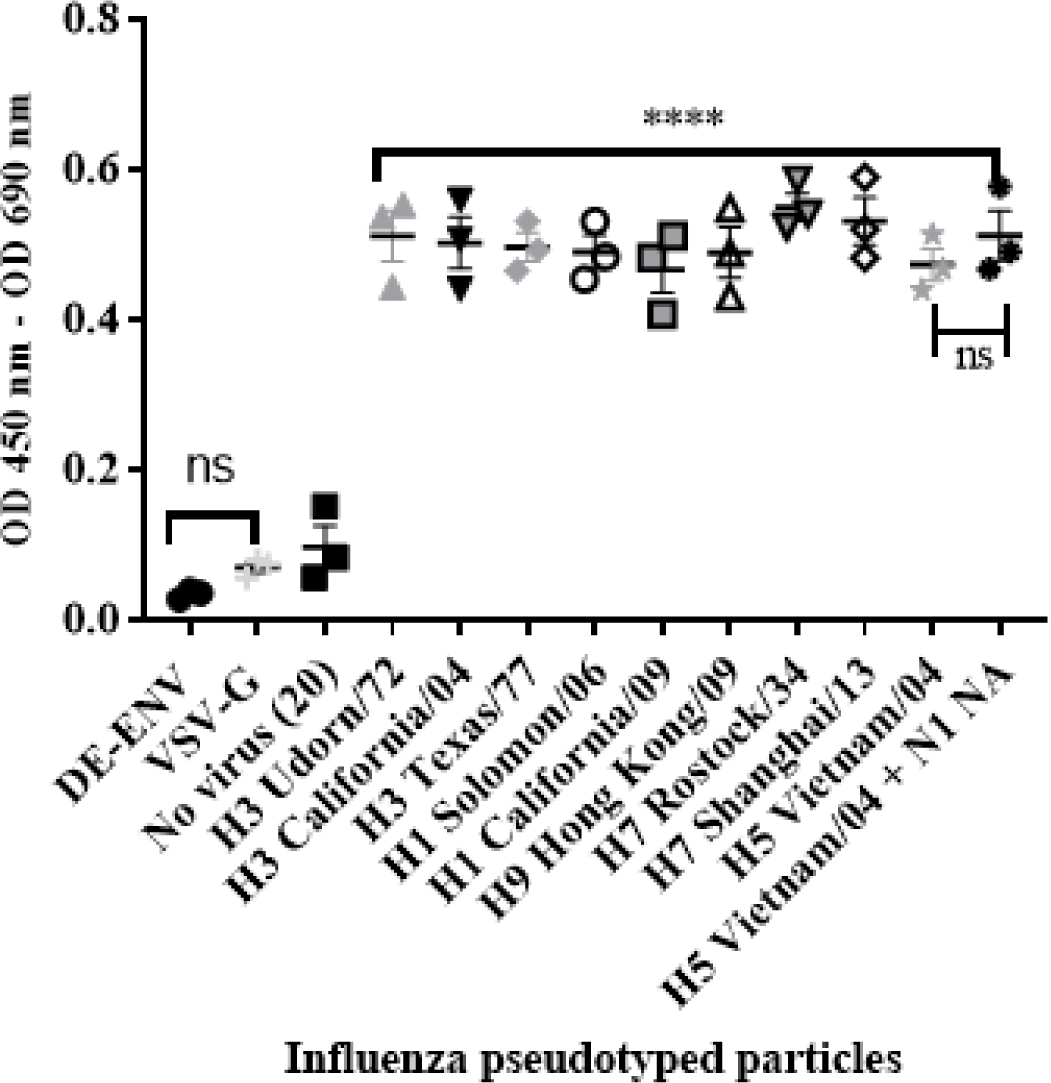
HA protein mediates interaction between human FH and IAV. (A)10^4^ RLU/ well of lentiviruses pseduotyped with various envelope proteins were used in an ELISA with 20 µg/well of purified FH protein alongside the controls of 20 µg/well FH with no virus [No virus (20)], lentivirus particles with VSV-G in the envelope (VSV-G) and lentivirus particles generated with no envelope protein (DE-ENV). FH binding was detected by anti-FH monoclonal antibody (OX24). Mean OD 450nm – OD 690nm values with three independent experiments carrying three replicates per experiment are displayed with error bars indicating standard error of the mean. Significant binding was calculated by One-way ANOVA and post hoc Dunnetts multiple comparison test to the control group of No virus (20); **** p ≤0.0001.

### FH binding footprint on HA is in the region of the receptor binding site

To map the footprint of FH binding to the H1N1 (A/England/195/09) and H3N2 (A/Hong Kong/1774/99) HA proteins, we used a panel of N-terminal biotinylated 20mer peptides from the HA proteins that overlapped by 10 amino acids with the preceding and following peptides (Supplementary Table 1 & 2). The peptides solubility was confirmed by dot blot using HRP conjugated Streptavidin (Supplementary Figure 2B and C). An ELISA was performed whereby the biotinylated peptides were immobilised on streptavidin-coated microtiter wells and the binding of FH to the peptides was examined using the anti-FH monoclonal antibody OX24 (Table 1; Figure 4). We identified the binding footprint of FH on both the H1 and H3 proteins. Similarly, for both HAs, FH bound to peptides that made up the receptor binding site, with both the amino acids in HA1 of the 130-loop and 220-loop binding, suggesting that binding of FH to HA might interfere with the ability to bind the virus receptor, i.e. sialic acid (Figure 4C- F). There were more peptides spanning the H3 HA protein that bound to FH compared to the H1 HA (Figure 4), with peptides which form the bottom of the fusion peptide pocket in HA2 of the H3 HA structure but not the H1 HA also binding to FH.

**Figure 4.**
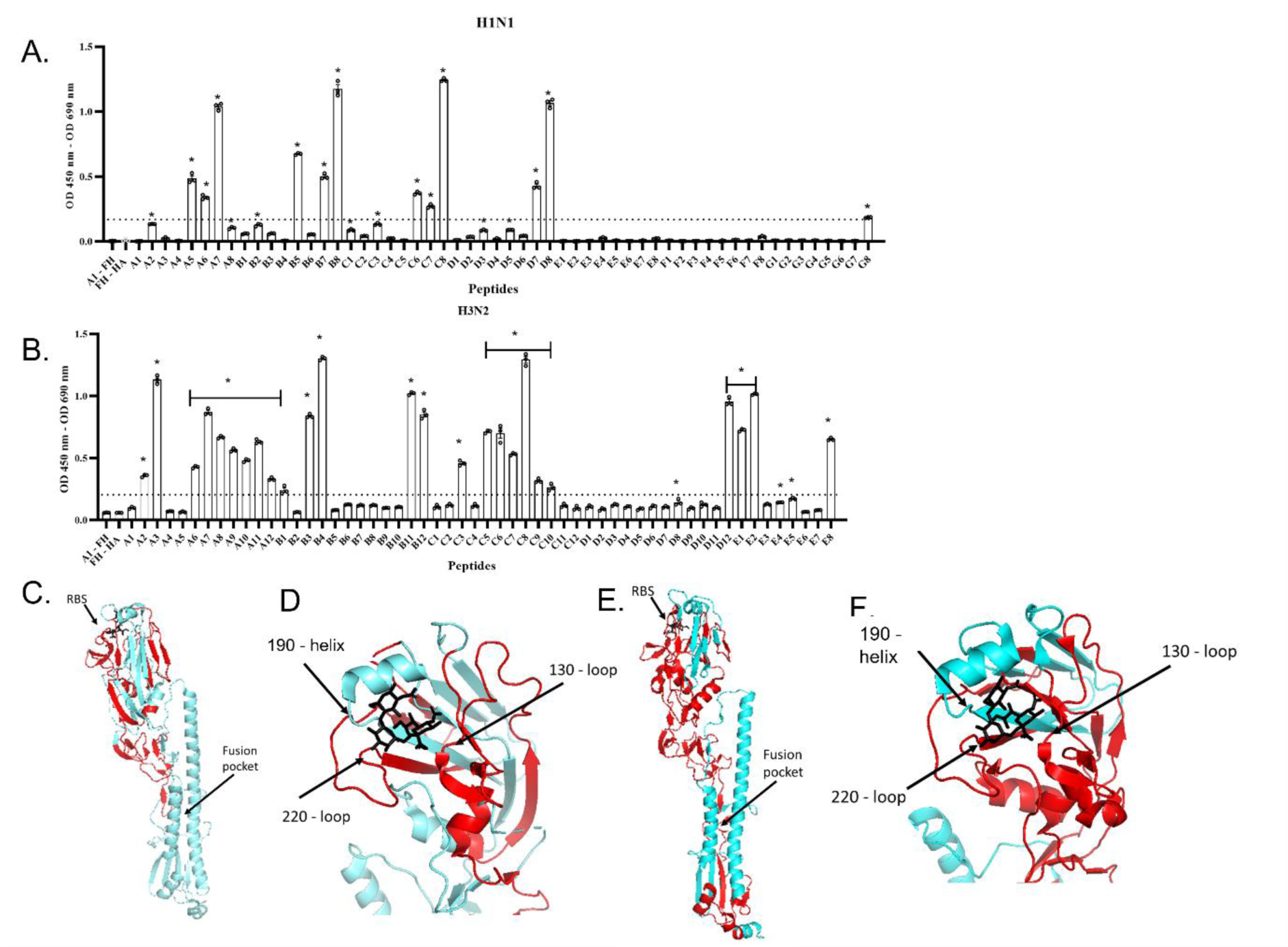
Human FH binds to the receptor binding region and fusion peptide pocket of IAV HA proteins. 5 µg/well of purified FH was used in ELISA assay with 2.5 µg/well of biotinylated 20-mer overlapping HA peptides. (A) H1N1 (A/England/195/09) and (B) H3N2 (A/Hong Kong/1774/99). Binding of the peptides to the FH is shown as mean OD 450nm – OD 690nm values are displayed with error bars indicating standard error of the mean. Negative controls whereby A1 peptide with no FH (A1 – FH) and FH with no HA peptide applied (FH – HA) were carried out alongside. Significant binding was calculated by One-way ANOVA and post hoc Dunnett’s multiple comparison test to the control group of FH – HA; * p ≤0.0001. Dashed line indicated mean difference from FH – HA of greater than 0.15 OD. The panel is representative of two independent experiments carrying three replicates per experiment. (C-D) H1 HA structure (A/California/7/09) (protein data bank 3UBN) (E-F) H3 HA structure (A/Finland/486/2004) (protein data bank 2YP3) indicating FH binding footprint (red), sialic acid (black) in the receptor binding site (RBS). Major structural features RBS and fusion peptide pocket are indicated and structural RBS loops 130, 180 and 220 marked. Images rendered in The PyMOL Molecular Graphics System, Version 2.0 Schrödinger, LLC.

### Human FH inhibits cell entry by human IAVs

The HA protein of IAVs contains the sialic acid receptor binding site that mediates viral attachment to host cells. In addition, HA possesses a pH activated fusion peptide which facilitates cell membrane fusion between the IAV virion and cellular endosomal membranes to permit genome uncoating. We hypothesised that binding of HA protein to human FH may affect the entry process of IAV during infection of cells. This hypothesis was tested by pre-incubating human IAVs with FH prior to a single round of infection in human lung cell line, A549 at an MOI of 1. At 4 and 8 hours post infection, the infected A549 cells were either fixed with ice cold 50:50 Methanol: acetone or lysed. Fixed cells were immune-stained with an anti-nucleoprotein (NP) antibody (Figures 5A and B) whereas cell lysates were separated by denaturing SDS-PAGE and transferred for western blotting and then probed with an anti-Matrix (M1) antibody (Figures 5C and D). At 4 hours post infection, we found that both human H1N1 (A/England/195/09) and H3N2 (A/Hong Kong/1774) IAV that had been preincubated with FH prior to infection showed reduced entry to A549 cells by both immune-staining and western blot analysis compared to IAV that had not been pre-incubated with FH (Figure 5A & C). The entry inhibition was greater for the human H3N2 strain compared to the H1N1 with almost a complete block noted in the case of H3N2. When we analysed the infection of the A459 cells at 8 hours post infection for H3N2, we again observed this near complete inhibition of entry (Figure 5B & D). Interestingly, for the H1N1 IAV strain, we observed an enhancement of the amount of viral protein in the cells at 8 hours post infection when the H1N1 virus was pre-incubated with FH protein (Figure 5B & D). For the cell loading control, we used β-tubulin, and this does not appear to change between the FH treated and untreated infected cells. Thus, the enhancement in IAV protein at 8 hours for H1N1 is not due to more cells being present in the samples.

**Figure 5.**
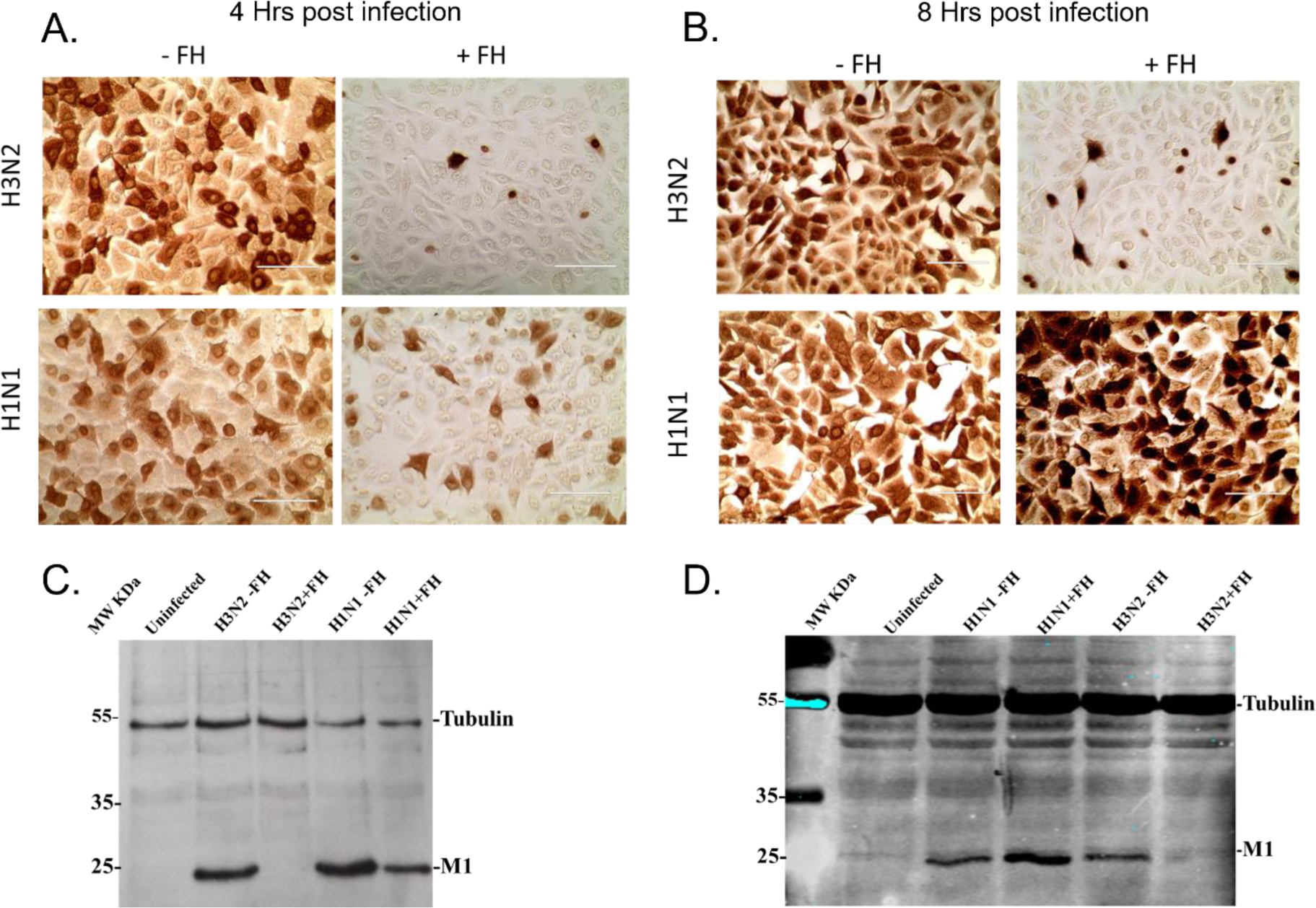
Pre-incubation of FH with human IAV strains alters virus entry to human lung cells. A549 cells were infected with a MOI 1 of H1N1 (A/England/195/09) or H3N2 (A/Hong Kong/1174/99) IAV that was preincubated with (+FH) or without FH (-FH). At 4 hours (A) or 8 hours post infection (B), cells were fixed and immune-stained with anti-NP antibody (brown staining). At 4 hours (C) or 8 hours post infection (D), cells were lysed and cell lysate was subjected to reducing SDS-PAGE. Proteins were western blotted, and the membrane probed using anti-M1 to detect IAV Matrix protein and anti-β tubulin as a loading control. Pre-stained MW protein marker was used as a reference (Thermo Scientific).

### Interaction of FH with H3N2 IAV inhibits replication in A549 cells

We used a low MOI multi-cycle growth assay to determine whether the interaction of FH with IAV strains which affected the entry of human IAV in A549 cells (Figure 6) altered the ability of the viruses to replicate and release progeny from these cells. The virus titres measured at 24 hours post infection demonstrated a strain specific alteration in virus replication (Figure 6). For the human strain H1N1 (A/England/195/09), there was no significant effect on titres collected from the A549 cells following preincubation of the virus with various concentrations of FH. In contrast the released human H3N2 (A/Hong Kong/1774/99) virus titre was significantly lower in all tested concentrations of FH preincubation. Both H9N2 (A/chicken/Pakistan/UDL01/08) and H5N3 (A/duck/Singapore/3/97) IAV titres were not affected by preincubation with 100 µg/ml of FH.

**Figure 6.**
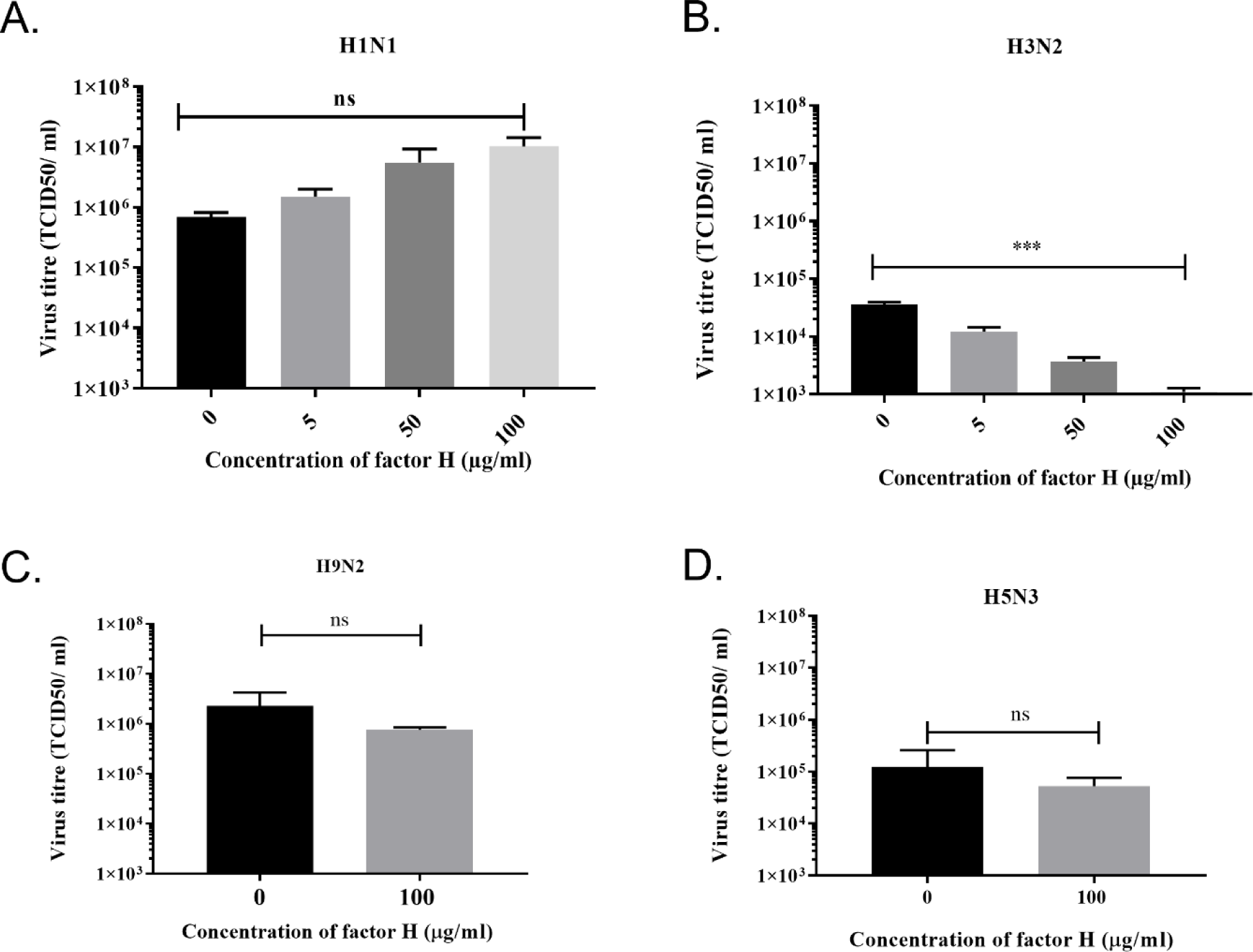
Preincubation of FH with H3N2 IAV inhibits replication in A549 cells. A549 cells were infected with an MOI 0.1 of (A) H1N1 (A/England/195/09) or (B) H3N2 (A/Hong Kong/1174/99) IAV that had been preincubated with various concentrations of FH (0, 5, 50 or 100 µg). A549 cells were infected with an MOI 0.1 of (C) H9N2 (A/chicken/Pakistan/UDL01/08) or (D) H5N3 (A/duck/Singapore/3/97 IAV that were preincubated with 100 µg of FH. 24 hours post infection, the cell supernatants were removed from infected A549 cells and titrated for infectious virus by TCID50 assay. The mean viral titre (TCID50/ml) is plotted in each panel with error bars indicating standard deviation. Unpaired, two-tailed t-test to 0 µg/ml FH was performed; *** p ≤0.001.

## Discussion

Factor H (FH) is a negative regulator of the complement alternative pathway. It functions to protect “self” cells from lysis by the complement system. Multiple pathogens including bacteria and viruses have the capacity to recruit FH to their surfaces as a complement evasion strategy [22]. In this study, we validated the earlier report that FH can bind to human IAVs of the H1N1 and H3N2 subtype [28]. In addition, we expand the observation to demonstrate that avian influenza viruses of the H9 and H5 subtypes are also able to bind human FH, suggesting that the interaction is conserved amongst IAV subtypes. FH is abundant in plasma at concentrations ranging from 116 µg - 562 µg/ml [22]; however, there is evidence that FH is present on mucosal surfaces and in the airway lumen as a result of production by epithelial cells and leakage from the plasma to these surfaces under inflammatory conditions and this concentration is around 10% of plasma levels (10µg - 50µg/ml) [30, 31]. This means that FH is available for interaction during natural IAV infection scenarios at the mucosal airway surfaces. The interactions we noted were significant at physiological concentration of FH, down to 50µg/ml (5µg/ well) (Figure 1) for most strains tested, H1N1 A/Michigan/45/2015 being the only exception which showed significant binding at 200µg/ml (20µg/well) (Figure 1B).

FH is composed of 20 complement control protein (CCP) modules (1-20), with CCPs 1-4 known to be involved in C3b binding, decay acceleration of C3 convertase and binding as a cofactor for factor I. Two other C3b binding sites have been mapped to CCP7-15 and CCP19-20, whereas the predominant binding to “self” cells via polyanions is via CCP19-20 [32, 33]. Interestingly, studies involving the mapping of FH-binding sites on other pathogens such as *Borrelia spp., Neisseria spp., Haemophilus influenzae, Pseudomonas aeruginosa, Candida albicans* and *Staphylococcus aureus* have implicated a common binding footprint for these diverse interaction molecules with either CCP5-7 and/ or CCP19-20. There are exceptions to this with the binding site for *Streptococcus pneumonia, Streptococcus agalactiae* and *Onchocerca volvulus* being outside these common regions. Using recombinant CCP fragments of FH, we mapped the binding footprint on FH for IAV strains and found that CCP5-7 and CCP15-20 mediated binding of four different IAV subtypes (Figure 2) and that IAV also bound to the common pathogen interaction sites on FH leaving the regulatory CCP1-4 available for complement inhibitory functions. Further work is required to narrow down the footprint precisely and determine if FH can still inhibit C3 when bound to IAV and thus act as a shield from the complement alternative pathway.

The surface of IAVs contain three viral proteins: haemagglutinin (HA), Neuraminidase (NA) and the ion channel M2. It has been shown previously, using far western blot under reducing conditions, that FH may interact with HA, NA, and the Matrix protein (M1) from IAVs [28]. Since HA is the most abundant protein on the surface of influenza viruses, we examined the HA derived from a range of human and avian IAV strains in the context of pseudotyped lentivirus particles and found that HA alone was able to bind to FH in an ELISA (Figure 3). The HA protein of IAV has multiple functions in the viral lifecycle, which include binding to the cellular receptor, sialic acid, to mediate attachment and insertion of the fusion peptide into the endosomal membrane to allow viral genome entry into the cell. In addition, it is the primary target of the antibody response of the host, and consequently, can be variable in sequence. Our mapping of the binding footprint of FH on the HA structure was carried out using overlapping peptides of the HA of both the H1 and H3 subtypes (Figure 4). The HA protein is comprised of two peptides (HA1 and HA2) held together by disulphide bonds. We found that FH predominantly bound to peptides located in the HA1 globular head of the HA protein which is the receptor binding site of IAV. The receptor binding site of HA is formed by two surface loops, the 220-loop (amino acids 225-228) and the 130- loop (amino acids 135-138), and the 190-helix (amino acids 187-194) [34]. For both the H1 and H3 HA, we found that FH could bind peptides that corresponded to all three sites that made up the receptor binding site, suggesting that FH may occlude the receptor binding site impeding attachment and entry of the virus to cells. Interestingly, there was some binding of FH to the membrane proximal stem region of HA for both the H1 and H3 peptide panel although this was more extensive for the H3 HA. The fusion peptide is comprised of the first 25 amino acids of the HA2 peptide (amino acids 330-355). The pocket into which the fusion peptide sits and initiates the conformational orientation change of the fusion peptide at low pH is made up of residues at the start of HA1 (amino acids in the region of 17) as well as HA2 residues (amino acids in the region of 380 and 440). FH bound to peptides that encompassed the fusion peptide and the fusion pocket in the H3 HA panel, suggesting likely impairment of the ability of the H3 to undergo the conformational change that leads to membrane fusion and release of viral genome into the host cell. However, the peptide library does not account for the natural state of the HA protein which forms a trimer, is inserted into the membrane of the virion and can be extensively glycosylated in a strain-specific manner, and therefore, the ability of FH to form these contacts with the natural HA protein as well as the strength of this interaction needs further investigation.

To assess whether the interaction of FH with the IAV HA affects life cycle functions of the virus, we carried out infection assays where IAVs were preincubated with FH before being allowed to infect a human cell line, A549s (Figure 5 and 6). We looked at the ability of the virus to attach to and enter the target cell at early time points and produce viral proteins (nucleoprotein, N or matrix protein, M) as well as the ability to complete a whole infectious lifecycle resulting in the release of infectious progenies from the infected cells. For both H1 and H3 IAV subtypes, we found virus entry was impeded at early time points, suggesting that the interaction with the receptor binding site in the HA does abrogate its ability to bind sialic acid. For the H3N2 subtype, there was a complete blockage of virus entry into cells that was not observed for the H1N1 subtype, possibly due to the double action of FH on the H3 HA of occluding the receptor binding site as well as binding to the fusion peptide preventing membrane fusion. In the presence of FH, even at the low airway lumen physiological concentration of 5 µg/ml, H3N2 virus was significantly inhibited (Figure 6). In contrast, for the H1N1 subtype, after the initial inhibition of virus entry caused by preincubation with FH, the virus replication cycle recovered to show robust infection and release of virus in A549 cells. This suggests that the interaction with the H1 subtype was either of lower affinity and competed off the HA more easily, or the lack of FH binding to the fusion peptide meant that once attachment had successfully occurred the FH posed no impediment to membrane fusion and viral genome release. FH bound to the avian IAVs tested but showed no impairment in full virus lifecycle replication in A549 cells at the concentration as high as 100 µg/ml of FH.

In summary, we show that FH can bind diverse IAV strains, including those isolated from avian hosts, and this binding is predominantly mediated by interaction with the viral HA protein. We have mapped the interaction footprint and shown that IAV binds to common CCP domains in human FH that other pathogens also bind to evade complement. In addition, the binding to the HA protein by FH is in the region of the receptor binding site but additional binding to the fusion peptide is strain specific. As a result of the strain specific interaction profiles of FH with HA proteins, the impact on viral replication is also strain specific. Further investigation is required to determine if the binding of FH to IAV is an impediment to the virus or, as seems to be the case for H1, H5 and H9 IAV subtypes tested here, there is no restriction to the virus. In the environment of the respiratory tract, recruitment of locally produced FH can potentially aid the survival of IAV by acting as a complement evasion strategy.

## Funding Information

This work described herein was funded by the Iraqi Ministry of Higher Education and Scientific Research and the University of Basrah, Iraq PhD studentship scholarship to Iman Rabeeah and BBSRC ISP Grant BBS/E/I/00007038 & BBS/E/I/00007034 to the Pirbright Institute. UK acknowledges funding via UAEU UPAR grant. The funders had no role in study design, data collection, data interpretation, or the decision to submit the work for publication.

## Acknowledgements

We would like to acknowledge members of the Influenza Virus group at The Pirbright Institute who supported this work; Dr Dagmara Bialy, Dr Klaudia Chrzastek and Dr Khalid Zakaria.

---

**Supplementary Figure 1.**
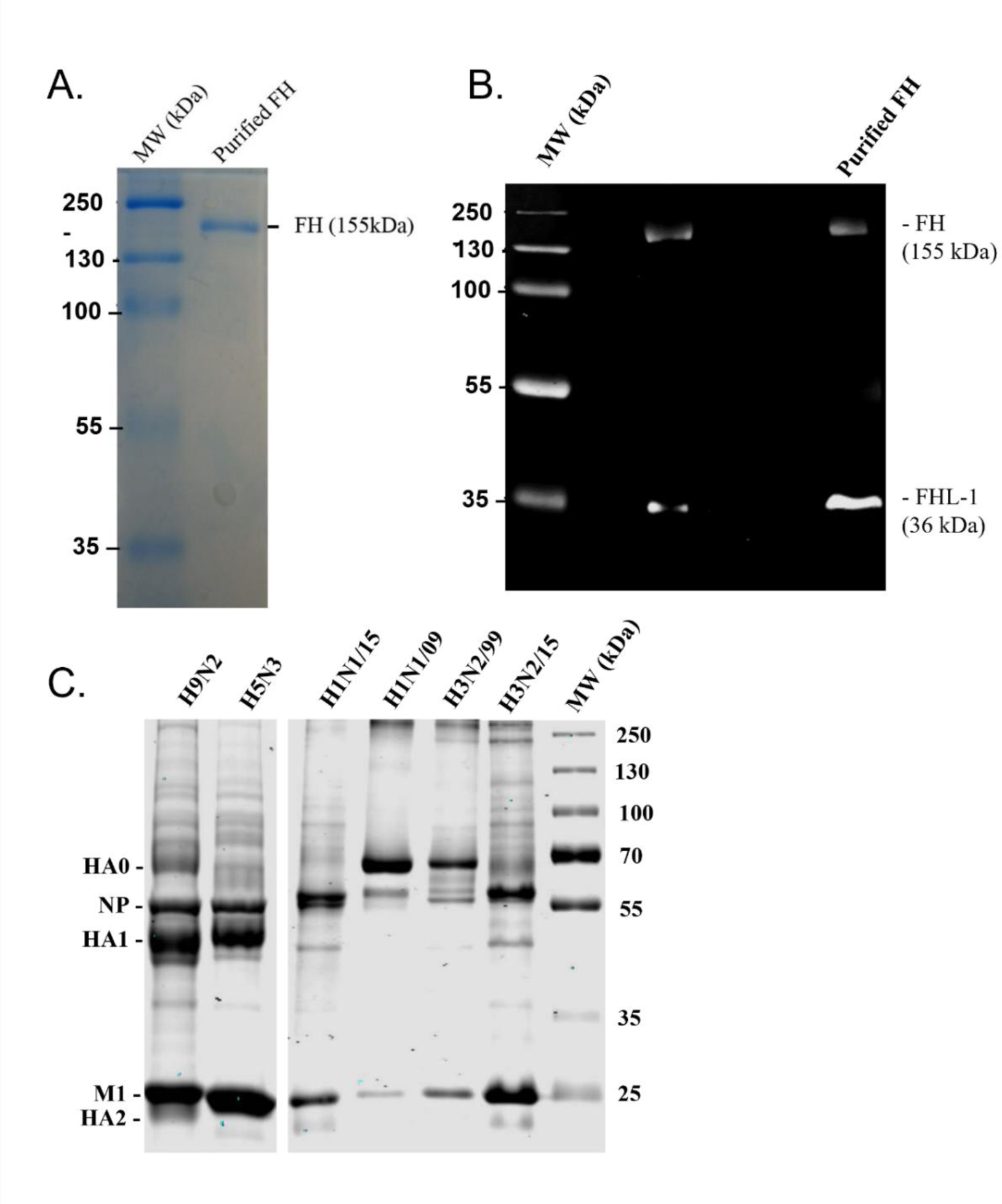
FH was purified from human plasma and subjected to non- reducing and reducing condition gel electrophoresis (A), followed by western blot with anti-FH OX24 antibody. Commercial purified FH (Com. FH) used as a control. (C) Coomassie-stained SDS-PAGE analysis of purified IAV preparations. The six strains of avian and human viruses in the panel were purified by sucrose gradient centrifugation and subjected to a 10% reducing SDS PAGE and stained with Coomassie blue. Lane 1: A/England/195/09 (H1N1pdm09), Lane 2: Vaccine strain, A/Michigan/45/15, (H1N1/15), Lane 3: the wild, H3N2 A/HK/1174/99, (H3N2/99), Lane 4: Vaccine strains A/HK/4801/14, (H3N2/14), Lane 5: Avian H5N3 A/Duck/Singapore/3/97, Lane 6: Avian H9N2 A/chicken/Pakistan/UDL-01/08, and Lane 7: the molecular size marker (MW). The locations of the viral proteins are characterised by their expected molecular weights.

**Supplementary Figure 2.**
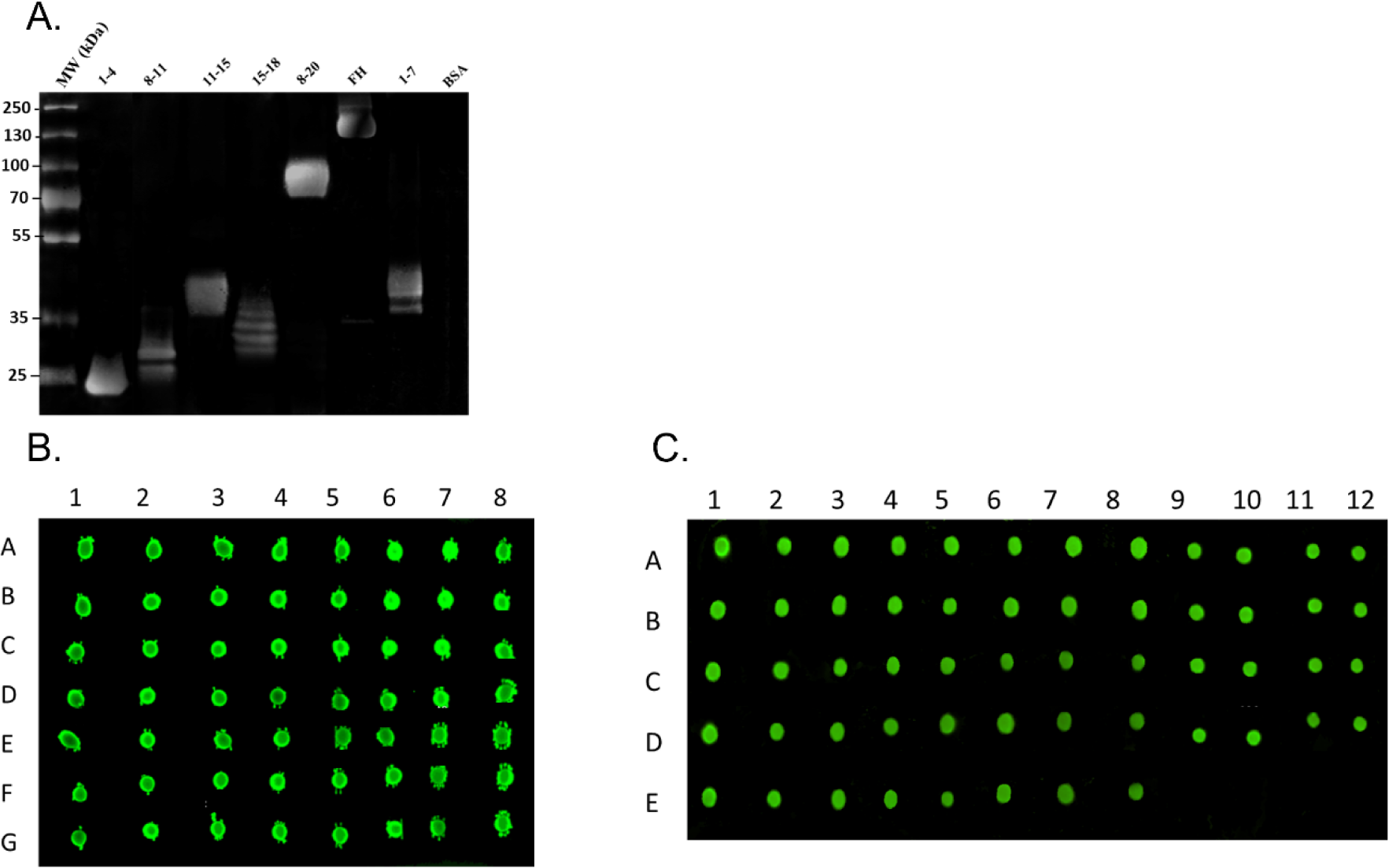
(A) Detection of FH fragments using anti-human FH antisera by western blot. 5 μg of each recombinant fragment - CCP1-4, CCP8-11, CCP11-15, CCP15-18, CCP8-20, and CCP1-7 were subjected to 10% reducing SDS- PAGE, followed by immunoblotting with polyclonal anti-human FH antiserum. Full- length FH was used as a positive control, whilst BSA was used as a negative control (5 μg each). The mobility of size markers is indicated to the left of the gel. Dot blot assay to confirm the solubility of synthetic HA peptides from (B) H1N1pdm09 and (C) H3N2/99. Each individual peptide (10 amino acids) was blotted on PVDF membrane with concentration of 2.5 μg. The membrane was then probed with IRDye 800CW HRP-conjugated Streptavidin at 1:500 dilution. As a negative control, BAS was used at the same dilution of HA peptide. The Odyssey Imaging System was used to visualise the interaction between the HA peptide and the HRP-conjugated streptavidin.

**Supplementary Table 1:**
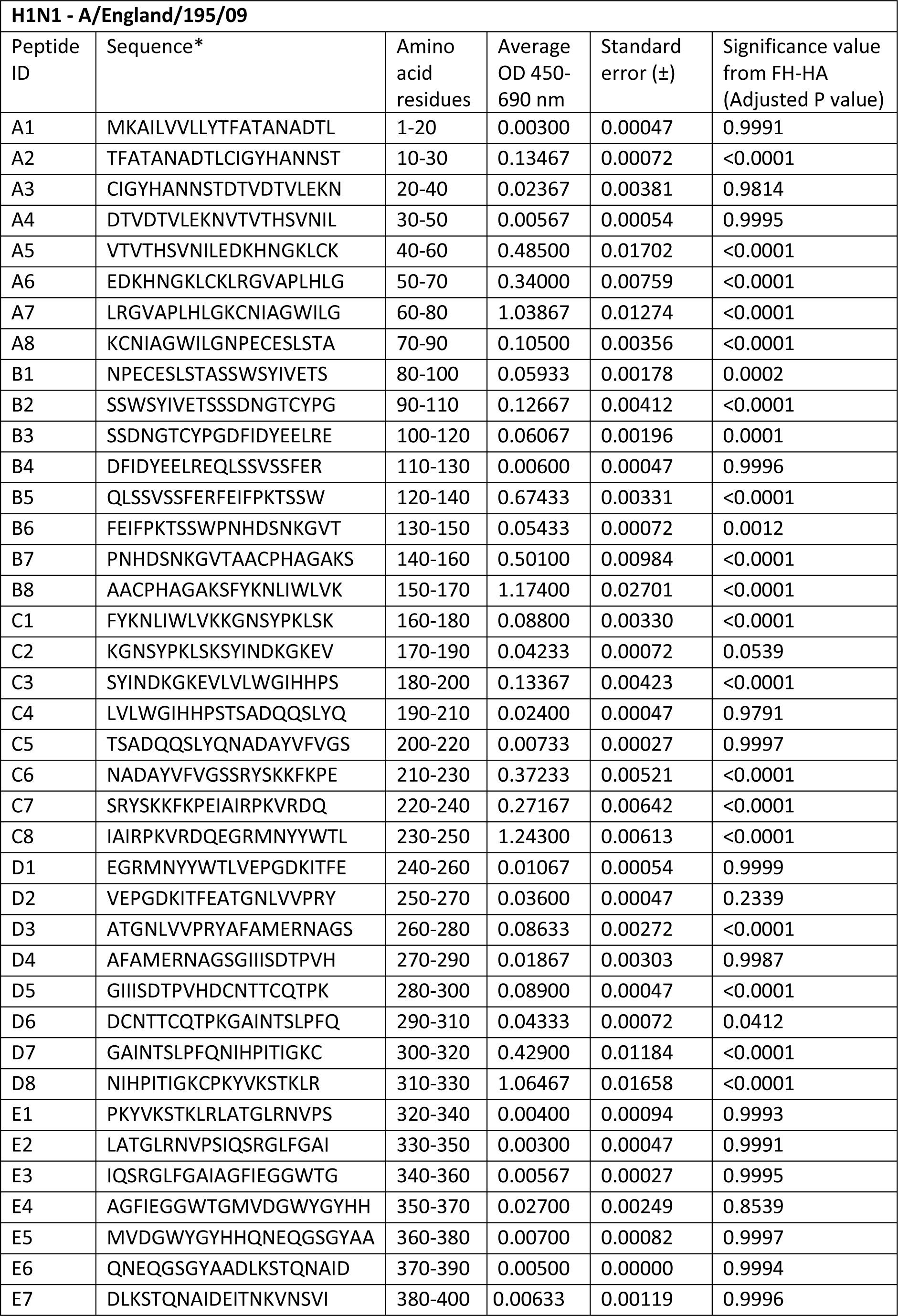
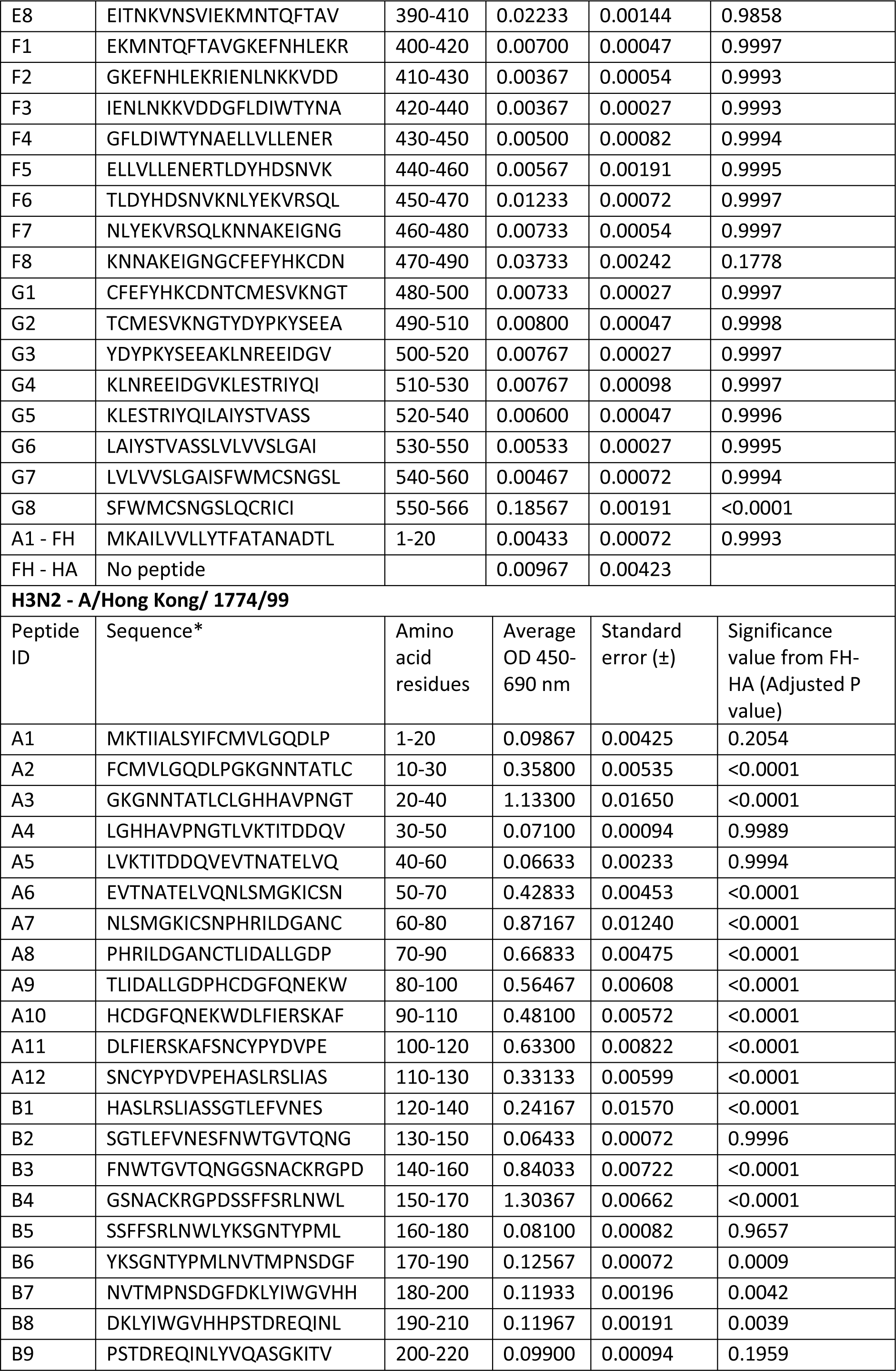
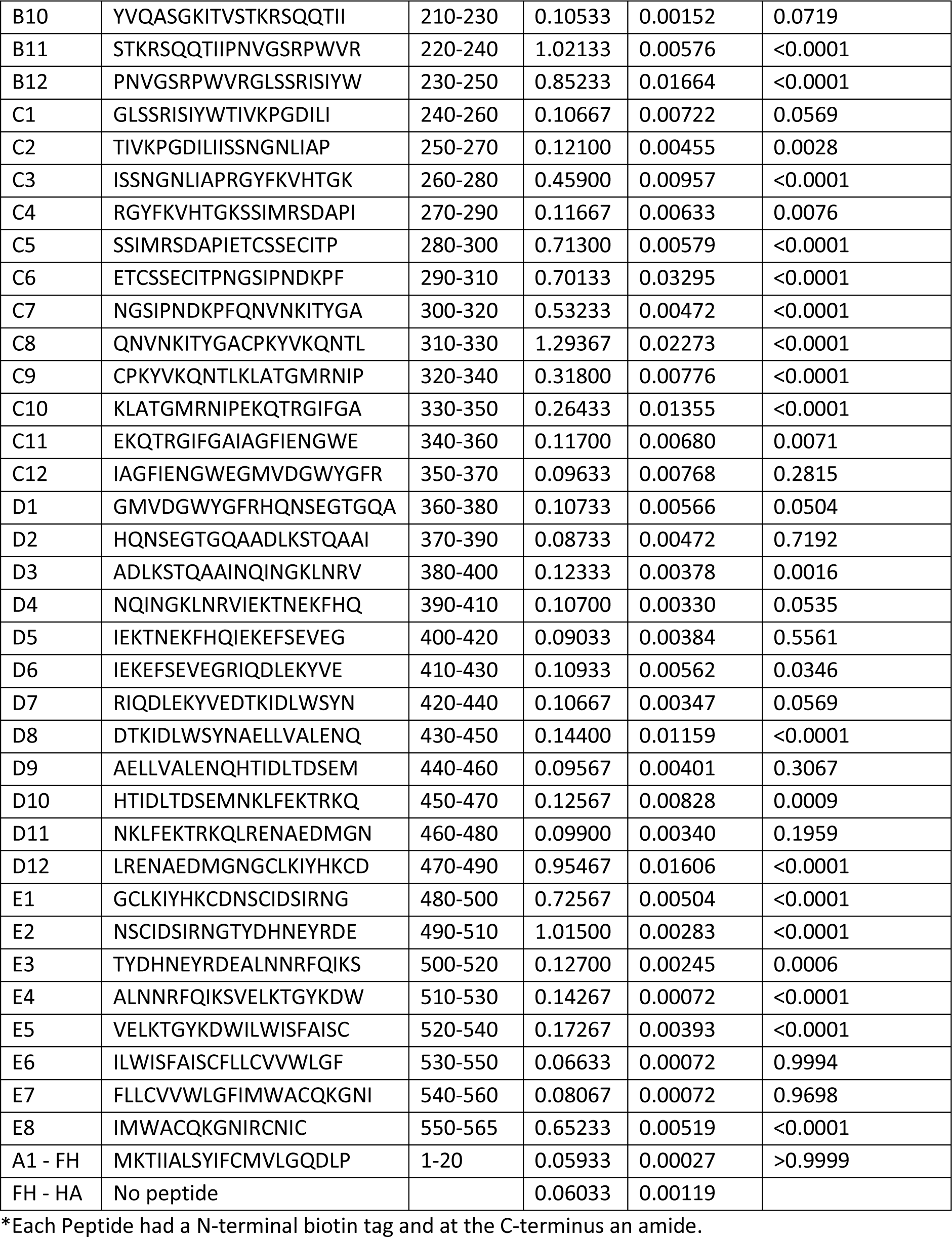
Sequence of H3N2 and H1N1 peptides used in this study.

## Notes

### Competing Interest Statement

The authors have declared no competing interest.

## References

[1] A.D. Iuliano, K.M. Roguski, H.H. Chang, D.J. Muscatello, R. Palekar, S. Tempia, C. Cohen, J.M. Gran, D. Schanzer, B.J. Cowling, P. Wu, J. Kyncl, L.W. Ang, M. Park, M. Redlberger-Fritz, H. Yu, L. Espenhain, A. Krishnan, G. Emukule, L. van Asten, S. Pereira da Silva, S. Aungkulanon, U. Buchholz, M.A. Widdowson, J.S. Bresee, N. Global Seasonal Influenza-associated Mortality Collaborator, Estimates of global seasonal influenza-associated respiratory mortality: a modelling study, Lancet 391(10127) (2018) 1285–1300.

[2] Z.Q. Wu, Y. Zhang, N. Zhao, Z. Yu, H. Pan, T.C. Chan, Z.R. Zhang, S.L. Liu, Comparative Epidemiology of Human Fatal Infections with Novel, High (H5N6 and H5N1) and Low (H7N9 and H9N2) Pathogenicity Avian Influenza A Viruses, Int J Environ Res Public Health 14(3) (2017).

[3] D.S.C. Hui, N. Lee, P.K.S. Chan, Avian influenza A (H7N9) virus infections in humans across five epidemics in mainland China, 2013-2017, J Thorac Dis 9(12) (2017) 4808–4811.

[4] Y.Y. Shen, C.W. Ke, Q. Li, R.Y. Yuan, D. Xiang, W.X. Jia, Y.D. Yu, L. Liu, C. Huang, W.B. Qi, R. Sikkema, J. Wu, M. Koopmans, M. Liao, Novel Reassortant Avian Influenza A(H5N6) Viruses in Humans, Guangdong, China, 2015, Emerg Infect Dis 22(8) (2016) 1507–9.

[5] N.S. Merle, S.E. Church, V. Fremeaux-Bacchi, L.T. Roumenina, Complement System Part I - Molecular Mechanisms of Activation and Regulation, Front Immunol 6 (2015) 262.

[6] N.S. Merle, R. Noe, L. Halbwachs-Mecarelli, V. Fremeaux-Bacchi, L.T. Roumenina, Complement System Part II: Role in Immunity, Front Immunol 6 (2015) 257.

[7] D.P. Beebe, R.D. Schreiber, N.R. Cooper, Neutralization of influenza virus by normal human sera: mechanisms involving antibody and complement, J Immunol 130(3) (1983) 1317–22.

[8] J.P. Jayasekera, E.A. Moseman, M.C. Carroll, Natural antibody and complement mediate neutralization of influenza virus in the absence of prior immunity, J Virol 81(7) (2007) 3487–94.

[9] J.T. Hicks, F.A. Ennis, E. Kim, M. Verbonitz, The importance of an intact complement pathway in recovery from a primary viral infection: influenza in decomplemented and in C5-deficient mice, J Immunol 121(4) (1978) 1437–45.

[10] M. Kopf, B. Abel, A. Gallimore, M. Carroll, M.F. Bachmann, Complement component C3 promotes T-cell priming and lung migration to control acute influenza virus infection, Nat Med 8(4) (2002) 373–8.

[11] E.M. Anders, C.A. Hartley, P.C. Reading, R.A. Ezekowitz, Complement-dependent neutralization of influenza virus by a serum mannose-binding lectin, J Gen Virol 75 (Pt 3) (1994) 615–22.

[12] Q. Pan, H. Chen, F. Wang, V.T. Jeza, W. Hou, Y. Zhao, T. Xiang, Y. Zhu, Y. Endo, T. Fujita, X.L. Zhang, L-ficolin binds to the glycoproteins hemagglutinin and neuraminidase and inhibits influenza A virus infection both in vitro and in vivo, J Innate Immun 4(3) (2012) 312–24.

[13] P. Agrawal, R. Nawadkar, H. Ojha, J. Kumar, A. Sahu, Complement Evasion Strategies of Viruses: An Overview, Frontiers in Microbiology 8(1117) (2017).

[14] M. Saifuddin, C.J. Parker, M.E. Peeples, M.K. Gorny, S. Zolla-Pazner, M. Ghassemi, I.A. Rooney, J.P. Atkinson, G.T. Spear, Role of virion-associated glycosylphosphatidylinositol-linked proteins CD55 and CD59 in complement resistance of cell line-derived and primary isolates of HIV-1, J Exp Med 182(2) (1995) 501–9.

[15] Y. Hidaka, Y. Sakai, Y. Toh, R. Mori, Glycoprotein C of herpes simplex virus type 1 is essential for the virus to evade antibody-independent complement-mediated virus inactivation and lysis of virus-infected cells, J Gen Virol 72 (Pt 4) (1991) 915–21.

[16] H.M. Friedman, L. Wang, N.O. Fishman, J.D. Lambris, R.J. Eisenberg, G.H. Cohen, J. Lubinski, Immune evasion properties of herpes simplex virus type 1 glycoprotein gC, J Virol 70(7) (1996) 4253–60.

[17] S. Mawatari, H. Uto, A. Ido, K. Nakashima, T. Suzuki, S. Kanmura, K. Kumagai, K. Oda, K. Tabu, T. Tamai, A. Moriuchi, M. Oketani, Y. Shimada, M. Sudoh, I. Shoji, H. Tsubouchi, Hepatitis C virus NS3/4A protease inhibits complement activation by cleaving complement component 4, PLoS One 8(12) (2013) e82094.

[18] B. Mazumdar, H. Kim, K. Meyer, S.K. Bose, A.M. Di Bisceglie, R.B. Ray, R. Ray, Hepatitis C virus proteins inhibit C3 complement production, J Virol 86(4) (2012) 2221–8.

[19] H. Kim, K. Meyer, A.M. Di Bisceglie, R. Ray, Hepatitis C virus suppresses C9 complement synthesis and impairs membrane attack complex function, J Virol 87(10) (2013) 5858–67.

[20] M.L. Shaw, K.L. Stone, C.M. Colangelo, E.E. Gulcicek, P. Palese, Cellular Proteins in Influenza Virus Particles, PLOS Pathogens 4(6) (2008) e1000085.

[21] J. Zhang, G. Li, X. Liu, Z. Wang, W. Liu, X. Ye, Influenza A virus M1 blocks the classical complement pathway through interacting with C1qA, J Gen Virol 90(Pt 11) (2009) 2751–2758.

[22] V.P. Ferreira, M.K. Pangburn, C. Cortés, Complement control protein factor H: the good, the bad, and the inadequate, Mol Immunol 47(13) (2010) 2187–97.

[23] C.Q. Schmidt, J.D. Lambris, D. Ricklin, Protection of host cells by complement regulators, Immunol Rev 274(1) (2016) 152–171.

[24] M. Józsi, Factor H Family Proteins in Complement Evasion of Microorganisms, Frontiers in Immunology 8(571) (2017).

[25] P.F. Zipfel, T. Hallstrom, S. Hammerschmidt, C. Skerka, The complement fitness factor H: role in human diseases and for immune escape of pathogens, like pneumococci, Vaccine 26 Suppl 8 (2008) I67–74.

[26] K.M. Chung, M.K. Liszewski, G. Nybakken, A.E. Davis, R.R. Townsend, D.H. Fremont, J.P. Atkinson, M.S. Diamond, West Nile virus nonstructural protein NS1 inhibits complement activation by binding the regulatory protein factor H, Proc Natl Acad Sci U S A 103(50) (2006) 19111–6.

[27] A.K. Zaiss, M.J. Cotter, L.R. White, S.A. Clark, N.C. Wong, V.M. Holers, J.S. Bartlett, D.A. Muruve, Complement is an essential component of the immune response to adeno-associated virus vectors, J Virol 82(6) (2008) 2727–40.

[28] V. Murugaiah, P.M. Varghese, S.M. Saleh, A.G. Tsolaki, S.H. Alrokayan, H.A. Khan, K.S. Collison, R.B. Sim, B. Nal, F.A. Al-Mohanna, U. Kishore, Complement-Independent Modulation of Influenza A Virus Infection by Factor H, Front Immunol 11 (2020) 355.

[29] S.R. Moore, S.S. Menon, C. Cortes, V.P. Ferreira, Hijacking Factor H for Complement Immune Evasion, Frontiers in Immunology 12(188) (2021).

[30] T. Sakaue, K. Takeuchi, T. Maeda, Y. Yamamoto, K. Nishi, I. Ohkubo, Factor H in porcine seminal plasma protects sperm against complement attack in genital tracts, J Biol Chem 285(3) (2010) 2184–92.

[31] C.G. Persson, J.S. Erjefält, L. Greiff, M. Andersson, I. Erjefält, R.W. Godfrey, M. Korsgren, M. Linden, F. Sundler, C. Svensson, Plasma-derived proteins in airway defence, disease and repair of epithelial injury, Eur Respir J 11(4) (1998) 958–70.

[32] T.S. Jokiranta, J. Hellwage, V. Koistinen, P.F. Zipfel, S. Meri, Each of the three binding sites on complement factor H interacts with a distinct site on C3b, J Biol Chem 275(36) (2000) 27657–62.

[33] M. Jozsi, M. Oppermann, J.D. Lambris, P.F. Zipfel, The C-terminus of complement factor H is essential for host cell protection, Mol Immunol 44(10) (2007) 2697–706.

[34] Y. Ha, D.J. Stevens, J.J. Skehel, D.C. Wiley, X-ray structure of the hemagglutinin of a potential H3 avian progenitor of the 1968 Hong Kong pandemic influenza virus, Virology 309(2) (2003) 209–18.

[35] N. van Doremalen, H. Shelton, K.L. Roberts, I.M. Jones, R.J. Pickles, C.I. Thompson, W.S. Barclay, A single amino acid in the HA of pH1N1 2009 influenza virus affects cell tropism in human airway epithelium, but not transmission in ferrets, PLoS One 6(10) (2011) e25755.

[36] J. James, W. Howard, M. Iqbal, V.K. Nair, W.S. Barclay, H. Shelton, Influenza A virus PB1-F2 protein prolongs viral shedding in chickens lengthening the transmission window, J Gen Virol 97(10) (2016) 2516–2527.

[37] L.J. Reed, H. Muench, A simple method of estimating 50% end points, American Journal of Epidemiology 27(3) (1938) 493–497.

[38] J.M.M. Del Rosario, K.A.S. da Costa, B. Asbach, F. Ferrara, M. Ferrari, D.A. Wells, G.S. Mann, V.O. Ameh, C.T. Sabeta, A.C. Banyard, R. Kinsley, S.D. Scott, R. Wagner, J.L. Heeney, G.W. Carnell, N.J. Temperton, Exploiting Pan Influenza A and Pan Influenza B Pseudotype Libraries for Efficient Vaccine Antigen Selection, Vaccines 9(7) (2021) 741.

[39] N.J. Temperton, K. Hoschler, D. Major, C. Nicolson, R. Manvell, V.M. Hien, Q. Ha do, M. de Jong, M. Zambon, Y. Takeuchi, R.A. Weiss, A sensitive retroviral pseudotype assay for influenza H5N1-neutralizing antibodies, Influenza Other Respir Viruses 1(3) (2007) 105–12.

[40] B.-B. Yu, B.E. Moffatt, M. Fedorova, C.G.S. Villiers, J.N. Arnold, E. Du, A. Swinkels, M.C. Li, A. Ryan, R.B. Sim, Purification, Quantification, and Functional Analysis of Complement Factor H, in: M. Gadjeva (Ed.), The Complement System: Methods and Protocols, Humana Press, Totowa, NJ, 2014, pp. 207–223.

[41] E. Sim, M.S. Palmer, M. Puklavec, R.B. Sim, Monoclonal antibodies against the complement control protein factor H (beta 1 H), Biosci Rep 3(12) (1983) 1119–31.

